# Visual dynamics cues in learning complex physical interactions

**DOI:** 10.1101/662353

**Authors:** Christopher J. Hasson, Paneed F. Jalili

## Abstract

This study investigated the role of visual dynamics cues (VDCs) in learning to interact with a complex physical system. Manual gait training was used as an exemplary case, as it requires therapists to control the non-trivial locomotor dynamics of patients. A virtual analog was developed that allowed naïve subjects to manipulate the leg of a virtual stroke survivor (a virtual patient; VP) walking on a treadmill using a small robotic manipulandum. The task was to make the VP’s leg pass through early, mid, and late swing gait targets. One group of subjects (n = 17) started practice seeing the VP’s affected thigh and shank (i.e., VDCs); a second control group (n = 16) only saw the point-of-contact (VP ankle). It was hypothesized that, if seeing the VP’s leg provides beneficial dynamics information, the VDC group would have better task performance and generalization than controls. Results were not supportive. Both groups had similar task performance, and for the late swing gait target, a decrement in manipulative accuracy was observed when VDCs were removed in a generalization task. This suggests that when learning to manipulate complex dynamics, VDCs can create a dependency that negatively affects generalization if the visual context is changed.

## INTRODUCTION

The performance of daily life tasks requires humans to interact with both familiar and unfamiliar objects with a range of dynamic complexities. Such activities include opening a door, drinking tea, driving a car, or interacting with another person. It is thought that humans accomplish this in part by internally modeling the dynamics of the interactions. An internal dynamics model (or representation) relates motor commands to object behavior as detected by the body’s sensors. These models can be used to predict how an object (or another person) will respond to a motor input, and what motor commands are needed to elicit a desired interaction^1^.

Proprioception has an established and critical role in learning novel dynamics^2–4^. However, visual cues can provide information that is unavailable to proprioception, which may aid in the development and recall of internal models. The visual presentation of object features (e.g., size, shape, or configuration) can inform expectations about static and dynamic object behavior. Seeing the geometry of familiar tools allows humans to correctly brace against reaction forces^5^ and can improve generalization^6^. Sighted object features can assist with estimation of object stability characteristics, mass distribution, and material properties^7–9^, and may influence how dynamics are represented in the nervous system^10^. Although prior studies have shown that visual dynamics cues (VDCs) can benefit learning^5–10^, they have primarily used objects with relatively simple dynamics (e.g., a hammer, spring, or block). It is otherwise unclear whether the benefits would be similar for more complex physical interactions.

An archetypical complex physical interaction is with another person. As an exemplary case, manual gait training requires therapists to control the complex locomotor dynamics of patients. Following along with a patient’s swinging leg and applying forces to achieve a desired kinematic trajectory is not trivial. A therapist must compensate for numerous forces, including those related to linked-segmental dynamics (e.g., gravitational, inertial, Coriolis, and centrifugal forces), as well as those from a patient’s muscular contractions. A recent study used a virtual analog of manual gait training to show that healthy adults are capable of forming internal representations of pathologic locomotor dynamics while training a simulated patient^11^. However, this study did not manipulate the visual information provided to the trainers, i.e., they could always see the simulated patient’s leg during manipulation. Thus, it is unclear whether the VDCs had a significant impact on the learning outcomes.

Therefore, this study aimed to investigate the role of visual dynamics cues (VDCs) in learning to interact with a complex physical system in the form of a locomoting virtual patient (VP). Naïve subjects were asked to manipulate the leg of a virtual stroke survivor walking on a treadmill, which was modeled as an end-point impedance that subjects interacted with using a small robotic manipulandum^11^. Subjects were asked to apply forces to the VP’s affected leg to make the VP’s ankle pass through three gait targets, which would normalize the gait pattern with respect to the unaffected leg. One group of subjects (n = 17) started practice seeing the VP’s affected thigh and shank; a second group (n = 16) only saw the point-of-contact (the VP’s ankle). If seeing the VPs leg provides learning benefits (e.g., by informing expectations about object behavior and constraining the space of possible dynamics models), it was hypothesized that subjects who saw the VP’s leg during practice would show improved manipulative performance and generalization relative to those that did not (control group).

## METHODS

### Virtual Patient (VP) Model

The VP was modeled as an end-point impedance that subjects interacted with using a small robotic manipulandum (GeoMagic Touch, 3D Systems, Andover MA). Full details of the model, which was based on previously collected data on stroke survivors^12^, are reported in Hasson and Goodman^11^. Using this stroke survivor data, a reference VP kinematic trajectory was specified (Figure 1; red trajectory). The step pattern associated with this reference trajectory (representing the affected leg) and had a smaller step height and width compared to the unaffected leg. When subjects applied a force that made the manipulandum deviate from the reference trajectory, they experienced an impedance force ***F_IMP_*** as given by

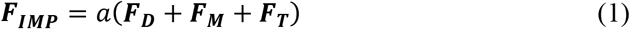

where the quantities ***F_D_***, ***F_M_***, and ***F_T_*** are vectors of forces that represent the anterior-poster and vertical components of impedance forces due to rigid body dynamics (***F_D_***), neuromuscular contributions (***F_M_***), and a treadmill interface (***F_T_***). The scaling factor *a* was determined through experimentation (*a* = 1/12) and was chosen so the amount of manipulative force applied by subjects to move the VP into a “healthy” trajectory was large enough to be significant without leading to fatigue (1-2 N).

**Figure 1.**
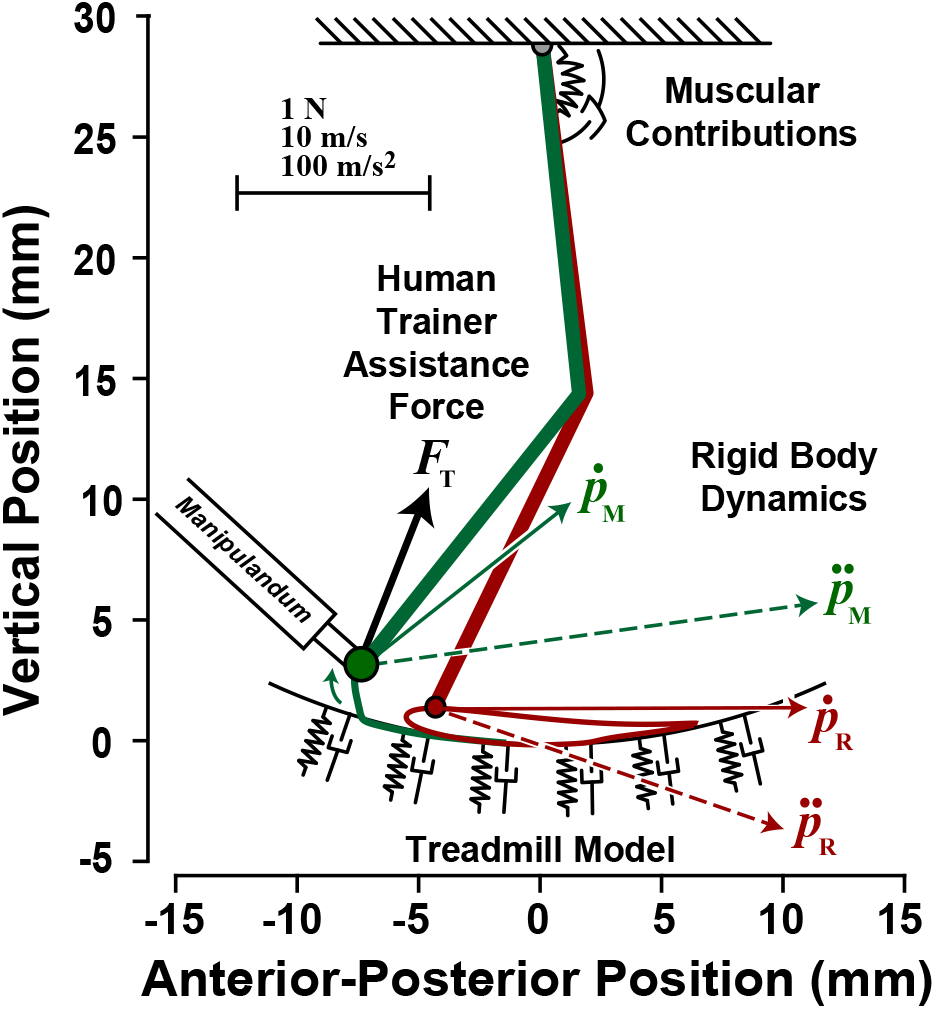
Schematic of virtual patient model. When a subject moved the manipulandum in a way that caused the virtual patient kinematics (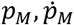 and 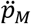, i.e., the virtual patient’s ankle position, velocity, and acceleration, respectively; green linked-segments and trajectory) to deviate from the reference stroke model kinematics (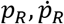, and 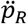; red trajectory), the subject felt an impedance force that depended on rigid body, muscular, and treadmill dynamics (the human trainer force is *F_T_*).

### Model Details

The rigid body forces (***F_D_***) that subjects felt when manipulating the VP’s leg were specified by

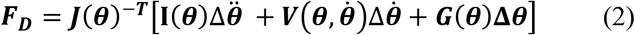

where **I**(***θ*)**, 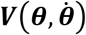 and ***G*(*θ*)** are matrices that specify inertial torques, centrifugal and Coriolis torques, and gravitational torques, respectively (details for **I**, ***V***, ***G*** are in Hasson and Goodman^11^). The quantities 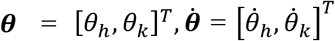, and 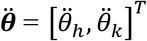 are vectors of joint angles, velocities, and accelerations, respectively. The subscripts *h* and *k* represent the VP hip and knee joints (superscript *T* denotes transposition). The quantities, Δ***θ***, 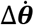, and 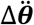 are difference vectors between the hip and knee angular kinematics of the reference stroke-survivor model and the kinematics of the VP as controlled by the subject (if the subject moves the manipulandum along the reference trajectory, these difference vectors are zero and the subject feels no rigid body kinetics). The rigid body torques are mapped to endpoint forces through multiplication with the inverse transpose of the Jacobian ***J*^−T^**, defined as

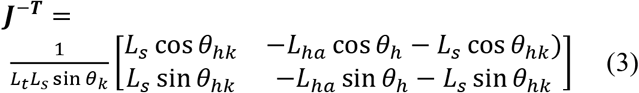

where *L_t_* and *L_s_* are the VP thigh and shank lengths, respectively, *L*_ha_ is the distance between the hip and ankle, and *θ_hk_*=*θ_h_*+*θ_k_*. The angular kinematics of the hip and knee can be similarly mapped to end-point kinematics for the reference model ankle joint position, velocity, and acceleration (*p_R_*, 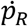, and 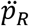, respectively; Figure 1) and the manipulandum (VP ankle) position, velocity, and acceleration (*p_M_*, 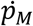, and 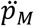, respectively; Figure 1).

Neuromuscular impedance forces (***F_M_***) depended on three quantities: a muscular spring stiffness ***K_SPR_***, a geometric stiffness ***K_GEO_,*** and a damping factor ***B***. The quantity ***K_SPR_***(***θ***,***ψ_h_***) modeled the spring-like action of human musculotendon units, which was a function of the segmental geometry ***θ*** and a variable-stiffness rotational hip spring with stiffness ***ψ_h_***, which was previously estimated from real patient muscular moments ***M*** and joint kinematics during locomotion^11,12^. ***K_GEO_***(***θ, M***) modeled how the limb endpoint stiffness due to muscular action is affected by skeletal geometry with a contact force^13^, which was a function of ***M*** and ***θ***. Finally, damping ***B***(***K_SPR_***) simulated the viscous effects of velocity-dependent muscular properties^14^, which was proportional to ***K_SPR_***. These viscoelastic effects were expressed as

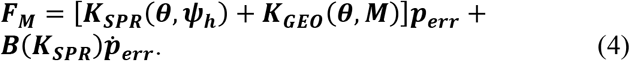

where ***p_err_*** and 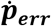 are difference vectors between the stroke reference and actual VP ankle (manipulandum) positions (***p_R_* – *p_M_***) and velocities 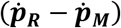, respectively (details for ***K_SPR_***, ***K_GEO_***, and ***B*** matrices can be found in Hasson and Goodman^11^).

Finally, a treadmill was created with a virtual spring (stiffness *k_N_* = 2.0 N/mm) that slid along a semicircular arc beneath the VP. A semicircular treadmill was needed because the VP’s leg was short one degree of freedom (the foot). An additional anti-slip spring (*k_S_* = 0.7 N/mm) prevented slipping due to frictional forces and dragged the VP leg rearwards during the stance phase. The net treadmill impedance force (***F_T_***) was given by

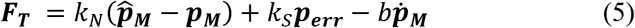

where 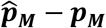 is a penetration vector, representing the difference between the actual manipulandum position ***p_M_*** and the projection of the manipulandum position on the treadmill surface 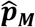, and *b* is a damping factor (*b* = 0.002), which increased fivefold (5*b*) at heel-strike for 0.1 s, linearly decreased to 1*b* over the next 0.1 s, and then remained at 1*b* until the next heel-strike. ***F_T_*** was set to zero if the ankle was above the treadmill surface.

### Subjects

Thirty-three young adults were randomly assigned to either a VDC group (n=17; 10 female, 7 male; age: 22 ± 1.7 yrs [mean ± SD]; height: 1.7 ± 0.1 m; weight: 67 ± 11 kg;) or a control group (n=16; 11 female, 5 male; age: 24 ± 4 yrs [mean ± SD]; height: 1.7 ± 0.1 m; weight: 67 ± 10 kg). The VDC group was able to see the VP’s leg during practice and the control group never saw the VP’s leg. All subjects were healthy and free from neurological and musculoskeletal impairments affecting upper extremity control and gave informed consent. The study was approved by the Northeastern University Institutional Review Board and all methods were performed in accordance with the relevant guidelines and regulations.

### Task Instructions & Goal

Subjects were told that they were interacting with a simulation of a stroke survivor (VP) walking on a treadmill and that if they push or pull on the manipulandum, the force is transferred to the VP’s ankle. Instructions were to make the VP’s ankle pass through three gait targets (Figure 2). These targets were: early, mid, and late swing, corresponding to 78, 88, and 100% of the VP’s gait cycle, respectively. These targets were selected from the average ankle trajectory of the unaffected leg for the stroke survivors upon which the model was based (hitting the targets increased VP gait symmetry).

**Figure 2.**
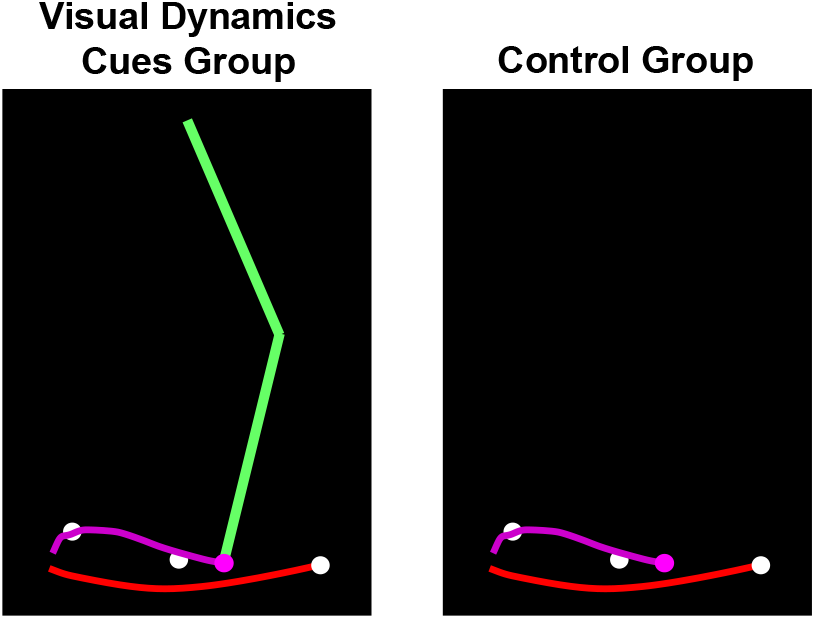
Illustration of visual feedback seen by study participants. The visual dynamics cues (VDC) group saw the affected leg of a virtual patient locomoting on a treadmill (left image) while a control group only saw the end-point (i.e., the ankle; right image). The white dots show early, mid, and late swing gait targets. For the VDC group, the leg segments are shown in green. For both groups, the end-point and trailing trajectory tail are shown in purple. The red curve at the bottom represents the path of the leg on the treadmill. The images show the affected leg during the swing phase after passing the mid-swing target, with the leg moving towards the right. The unaffected leg was not shown.

### Visual Display

All subjects saw the VP’s ankle (the interaction point), the three gait targets, a curved line representing the ground, and a magenta “tail” that followed the ankle to provide a visual indication of the ankle’s two-dimensional path (Figure 2). Importantly, the VDC group also saw two linked segments representing the VP’s thigh and shank. For both groups, a small rectangular tracker was presented when the VP’s leg was in contact with the treadmill to help subjects maintain proper step timing (because other timing-related visual information normally available during real gait training, such as a patient’s contralateral leg, was absent). The timing cue slid along the treadmill, becoming visible at heel-strike and disappearing at push-off. An audible beep was provided at heel-strike to assist with pacing. All programming associated with the virtual patient model and visual display was performed in C++ (Microsoft Visual Studio 2010 Professional, Version 10.0.30319.1; Microsoft Corporation, Redmond, WA).

### Initial Practice

Both groups began practice in a null-dynamics condition (two trials of 80 steps; Figure 3). The VDC group saw the VP’s leg and the control group did not. In this condition, subjects only experienced the treadmill model and felt no impedance forces from the VP (i.e. ***F_D_*** = ***F_M_*** = 0). The purpose of the null-dynamics condition was to allow subjects to become familiar with the task and interface. Next, subjects practiced the task with impaired-dynamics (Impaired Dynamics I). In this condition, subjects felt as if they were holding onto the leg of a miniature stroke patient (the VP) and had to apply appropriate forces to move the VP’s ankle through the three gait targets. In this condition, subjects performed four 80-step trials; the leg remained visible for the VDC group and absent for the control group. A one-minute rest was provided between each trial.

**Figure 3.**
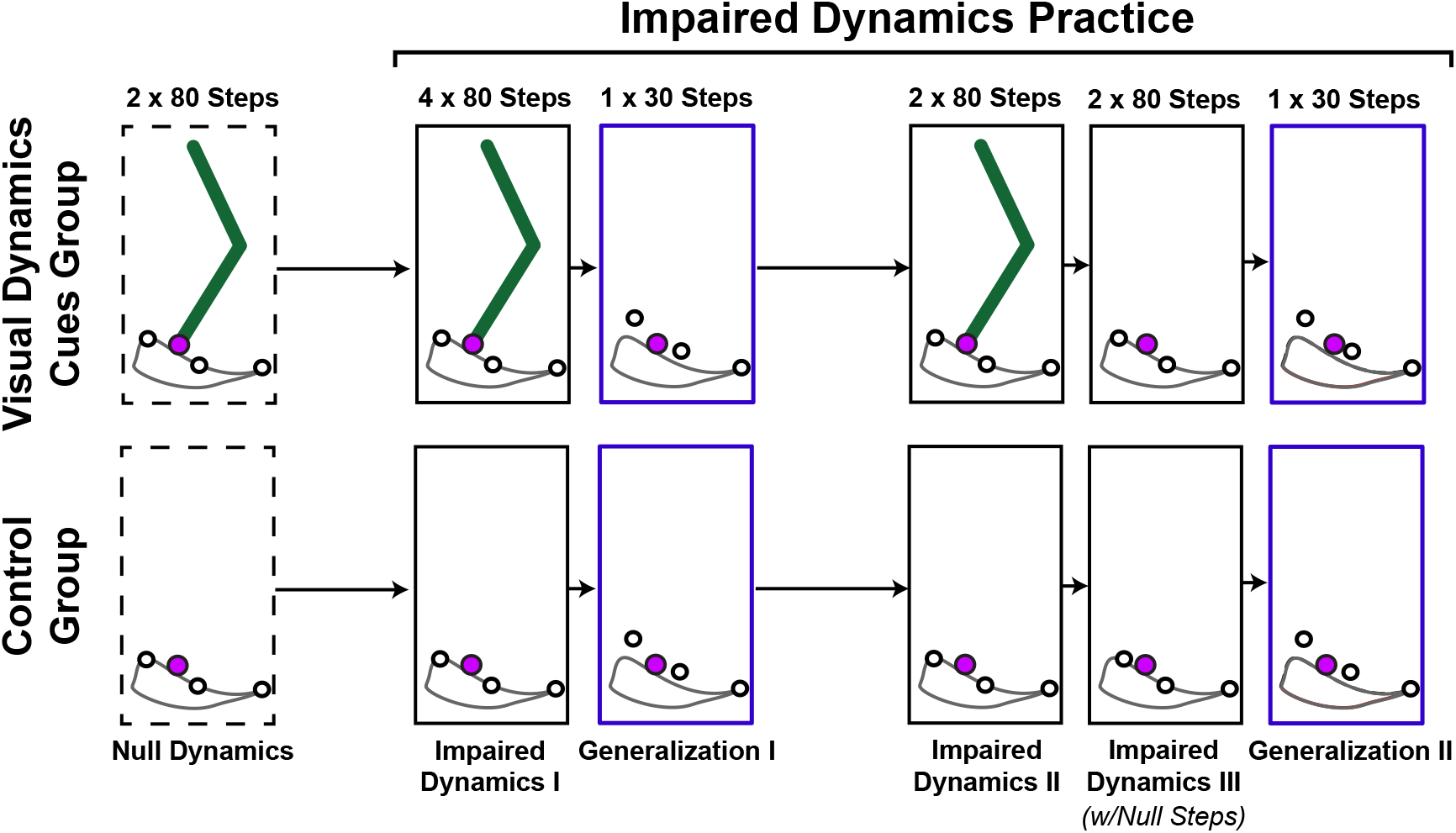
Experimental protocol for virtual locomotor manipulation task. One group began practice able to see the virtual patient’s thigh and shank segments (visual dynamics cues group); the other only saw the virtual patient’s ankle (the end-point).

### Generalization Tests

After initial practice, subjects performed two generalization tests. The early swing gait target was shifted 1.45 mm upwards (a 42% increase) and 0.24 mm to the right (see Figure 3). The mid-swing gait target height was increased by 1.21 mm. The late swing target was not changed. These target shifts required subjects to make the VP perform an exaggerated heel-lift during push-off. To facilitate group comparisons the generalization test was performed without the VP’s leg displayed so that both groups would have the same visual display. The first 30-step generalization test (Generalization I) was performed after Impaired Dynamics I. After more practice with the original targets (Impaired Dynamics II), a second 30-step generalization test was performed (Generalization II), but in this case, the VDC group had the opportunity to practice *without* seeing the leg before the target locations were changed (Impaired Dynamics III). This enabled the separation of two effects in the VDC group: one from the removal of the displayed leg segments and one due to switching target locations (Generalization I did both).

### Probe Steps

Errors made during unexpected changes in dynamics can reveal information about human sensorimotor control strategies^15^. Thus, each subject experienced four unexpected null-steps during the last practice trial before the second generalization test (Impaired Dynamics III; see Figure 3). In these steps, the VP rigid-body and neuromuscular impedance forces were turned off during the middle of the stance phase (the treadmill model remained in effect). Mid-stance was chosen as the initiation point because kinematic errors (and hence ***F_D_*** and ***F_M_*** impedance forces) are negligible at this time. The VP impedance forces remained off throughout the stance phase and were turned back on after the VP’s leg completed its swing (when the forward velocity of the VP’s ankle changed sign). The null-step numbers were selected from uniformly distributed pseudo-random integers in the span of 5-80 and were randomized within each subject; thus, null-step numbers differed across subjects. Each null-step had to be at least 4 steps from neighboring probe steps, if not a new random number draw was performed until the requirement was satisfied.

### Data Reduction and Analysis

The root-mean-squared error (RMSE) quantified task performance, determined as the smallest Euclidean distance between the manipulandum trajectory (the VP’s ankle) and the early, mid, and late swing gait targets (in two-dimensions). The RMSE was averaged across the first four steps of each practice condition to capture early practice performance, and across the last 20 steps to capture late practice performance (a shorter time scale was used for early practice because the changes are more rapid). For the null-trial analysis, the absolute RMSE and relative change in RMSE (ΔRMSE) were computed for each gait target and null-trial in the Impaired Dynamics III condition. For a given null-step, the ΔRMSE was calculated by subtracting the average RMSE during the four preceding regular (non-null field) steps from the RMSE on the null-step; thus, a positive ΔRMSE indicates the RMSE increased on the null-step. Since the null-step numbers were newly randomized for each subject (i.e., the null-step numbers differed across subjects), the absolute and relative null-step RMSEs were averaged across the four null-steps within each subject. For all analyses outliers were detected using Tukey fences^16^. If for a given condition and time point (early or late practice), a subject’s mean RMSE was outside ± 2.7 standard deviations (a 99.3% confidence interval), the value was excluded from further analysis (3.5% of data points). All analysis procedures were performed in MATLAB (ver. 9.4, R2018a; Mathworks, Natick MA).

### Statistics

The effects of VDCs cues on learning locomotor dynamics were assessed with a mixed models analysis. Group (VDC or control), Condition (Null Dynamics, Impaired Dynamics I, Generalization I, Impaired Dynamics II, Impaired Dynamics III, and Generalization II), and time (early vs. late practice within a condition) were included as fixed effects, with repeated measurements on condition and time (first-order autoregressive covariance structure). Subject ID was included as a random effect. The dependent variables were the RMSE for the early, mid, and late swing gait targets. The alpha level was .05 and all tests were two-tailed, which were performed in SPSS (ver. 25; IBM, Armonk, New York). The Benjamini-Hochberg (BH) step-up procedure^17^ was used to control for multiple comparisons with a tolerable false-discovery rate *Q* of 0.1^18^. BH critical values (*BH_CRIT_*) were calculated as *BH_CRIT_* = *Q*(*i*/*m*), where *m* is the number of tests (*m* = 12) and *i* is the p-value rank (smallest to largest). BH adjusted *p*-values are reported as “*q*” values, where *q* = *p*(*m*/*i*) and *p* is the raw *p*-value. Independent samples t-tests were performed to compare the absolute and relative null-trial RSME between the VDC and control groups during the Impaired Dynamics III condition for each gait target.

## RESULTS

Subject skill was assessed by the RMSE between the manipulandum (the VP’s ankle) trajectory and the early, mid, and late swing gait targets. For the early swing target both VDC and control groups reduced their error with practice (time main effect; F_1,151_ = 45.71, p < .001) with no effect of VDCs on RMSE (Figure 4; no group main effect; F_1,79_ = 0.50, p = .482). However, for the mid and late swing targets, there was a three-way interaction, indicating that there was an effect of visual cues on RMSE, but that the effect depended on the condition and time (mid-swing: F_5,204_ = 8.48, p < .001; late swing: F_5,280_ = 2.43, p = .047). Although three-way interactions can be difficult to interpret, the results in Figure 4 show that there was an effect of seeing the leg for the mid and late swing targets, but only for specific time points and conditions. This is explored further by analyzing pairwise differences.

**Figure 4.**
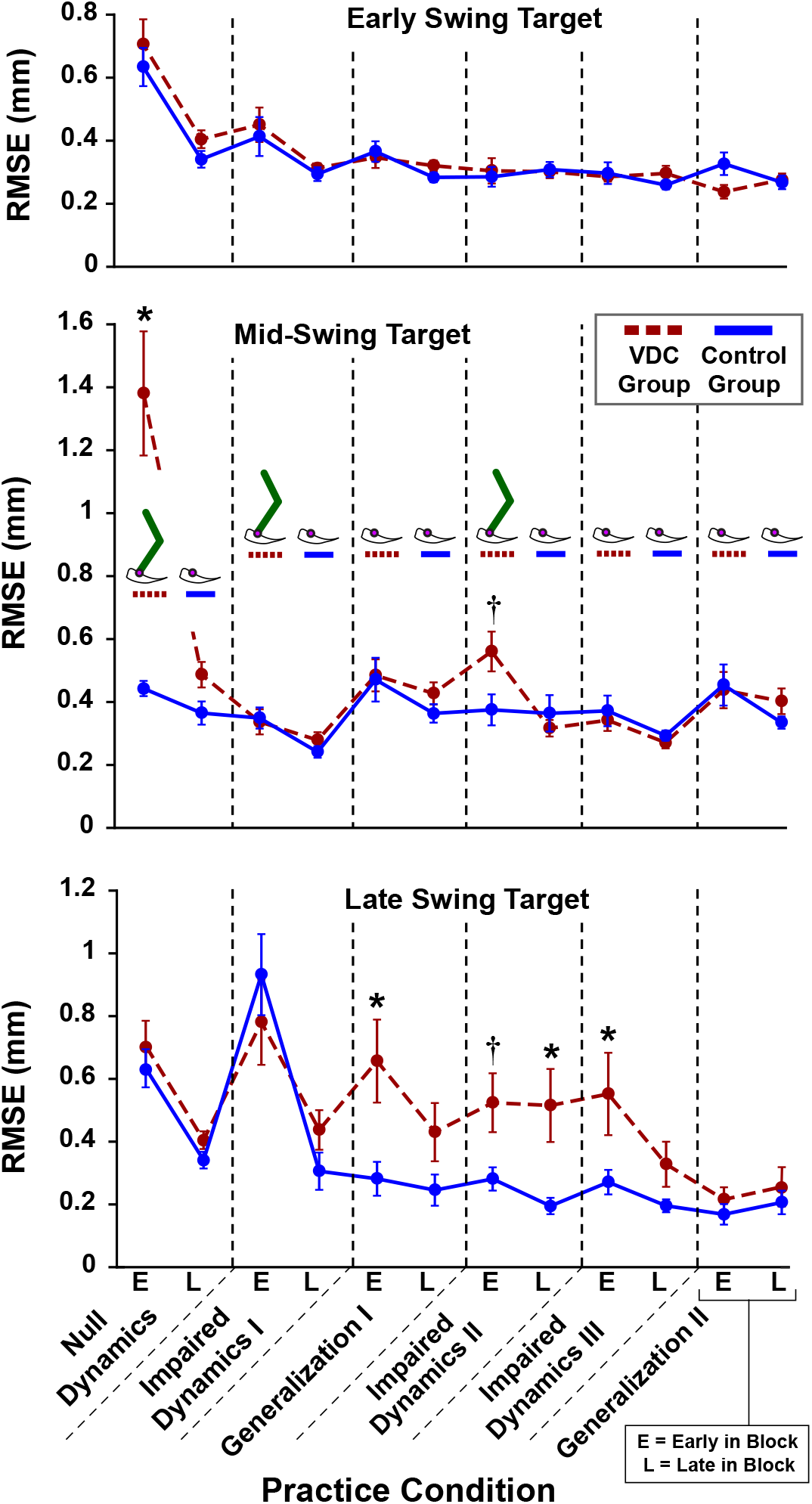
Task performance. Average root-mean-squared error (RMSE) for the early, mid, and late swing gait targets for the visual dynamics cues (VDC) group (red) and the control group (blue) while practicing the virtual locomotor training task. *Significant pairwise group comparisons after Benjamini-Hochberg procedure with false-discovery rate (FDR) of 0.1. †Non-significant after controlling FDR. Standard errors shown.

Pairwise comparisons showed that for the mid-swing target during early practice in Null Dynamics, the VDC group had larger errors than the control group (early practice *p* < .001 and *q* < .001), such that the VDC group significantly overshot the target (signed vertical error = 0.88 ± 0.82 mm; mean ± s.d.). Note that reported *q*-values are derived from the BH step-up procedure and are compared against the specified tolerable false-discovery rate (i.e., if *q* < 0.1 then the test remains significant). For the mid-swing target, both groups had a similar RMSE at the end of the Null Dynamics condition, and the group RMSEs remained similar throughout the rest of practice (the VDC group had a larger RMSE than controls during early practice in Impaired Dynamics II, but this was not significant after adjusting for multiple comparisons; *p* = .024 and *q* = .144). For the late swing target, the VDC group had a larger RMSE during early practice in Generalization I (*p* < .001 and *q* = .012), during late practice in Impaired Dynamics II (*p* = .015 and *q* = .090), and during early practice in Impaired Dynamics III (*p* = .016 and *q* = .064). By the end of Impaired Dynamics III, the group RMSE’s were similar (*p* = .094 and *q* = . 161).

To illustrate the nature of the errors made, representative scatterplots show the points of minimum RMSE, i.e., the closest approach of the VP’s ankle to the early, mid, and late swing gait targets in the Impaired Dynamics I and Generalization I conditions (Figure 5). Upon inspection, it is evident that each gait target had a distinct distribution of errors, with high asymmetry in the late swing target. The latter is due to the geometry of the VP’s leg and the constraints imposed on the VP, i.e., a fixed hip joint, hinged-rigid bodies that cannot pull apart (dislocate), and an arcing treadmill that compensated for the lack of a foot. Most errors for the late swing target reflected undershoots, in which the VP’s leg fell short and never reached the late swing target. Note that few points are seen past the late swing gait target because only the closest-approach is marked, so if the late swing target was passed, it is likely that the VP’s ankle would make a closer approach on its way back as the VP’s leg comes into contact with the virtual treadmill. The late swing target is where the greatest group differences were found, with the VDC group having larger undershoot errors when the targets were shifted in Generalization I (see Figure 4 and detail panels in Figure 5).

**Figure 5.**
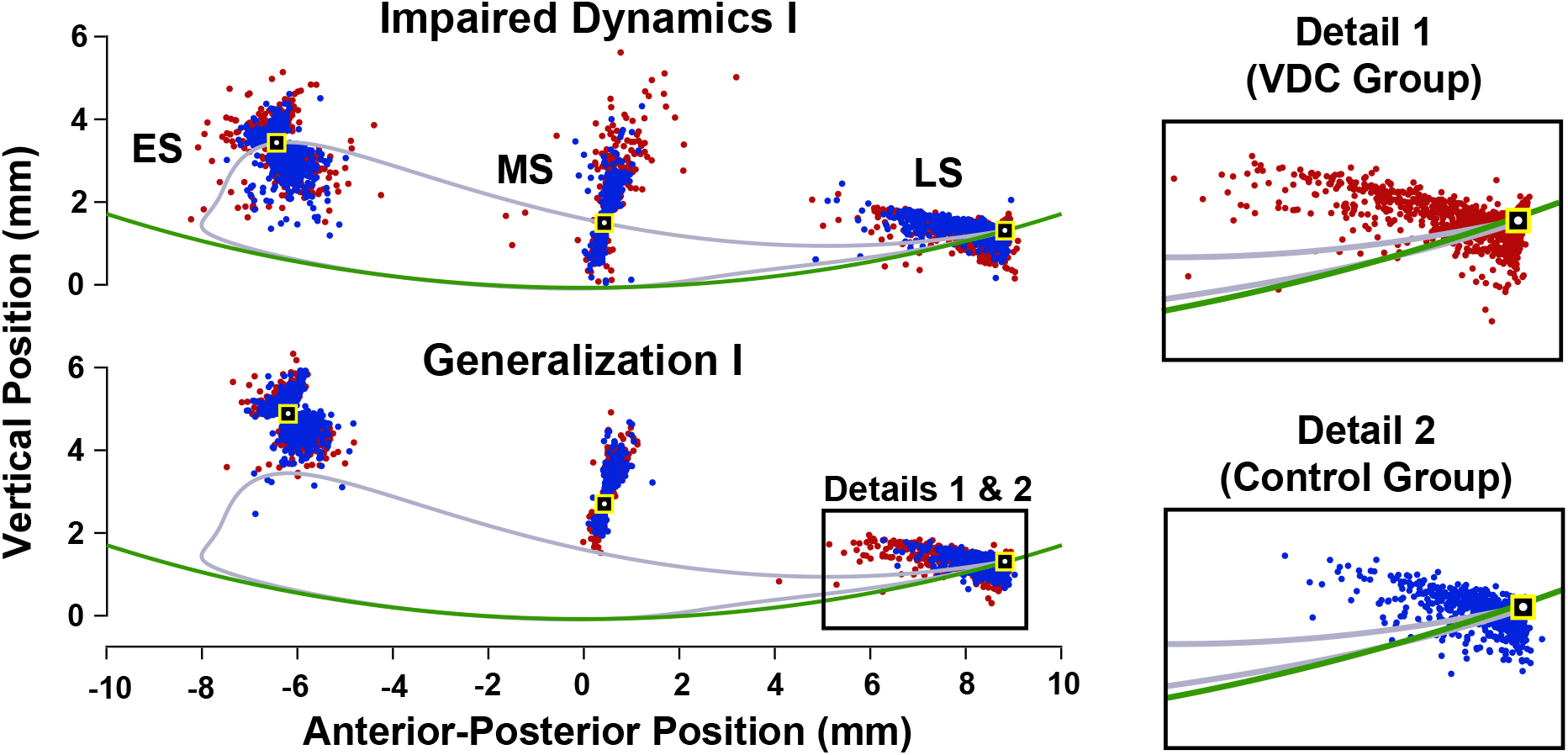
Detail of task performance showing error distributions. Each point represents the closest approach of the VP’s ankle (and manipulandum) to the early, mid, and late swing gait targets (ES, MS, and LS, respectively) for all steps and all subjects in the Impaired Dynamics I and Generalization I practice blocks. The red dots are from the visual dynamics cues (VDC) group and the blue dots are from the control group. The gray trajectory represents a fictitious “healthy” gait trajectory that was not shown to subjects. The gait targets, which subjects did see on the visual display, are shown as black and yellow squares with a white dot marking the center to enhance contrast. The green arc is the virtual treadmill surface.

Near the end of practice, during Impaired Dynamics III, subjects were exposed to four unexpected null-dynamics steps to probe the strength of participants’ internal representations. During these steps, subjects did not feel any impedance forces as they tried to lift the VP’s foot from the ground. As illustrated in Figure 6, the VDC group had a smaller absolute and relative RMSE for the early swing target during the null-step trials (absolute *p* = .037; relative *p* = .005). There were no group differences in null-trial RMSE for the mid-swing target (absolute *p* = .803; relative *p* = .948) and late swing target (absolute *p* = .702; relative *p* = .406).

**Figure 6.**
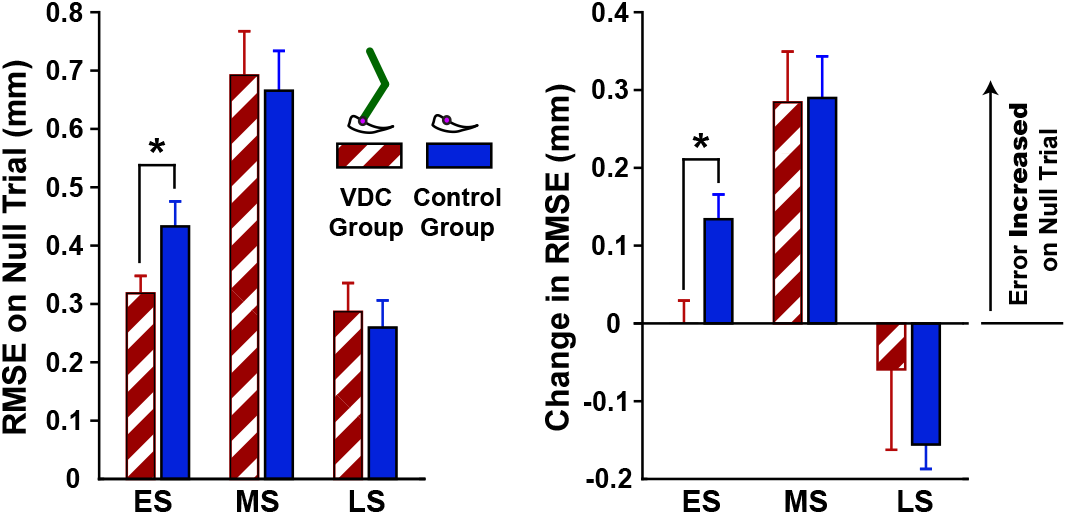
Null-step performance. The absolute root-mean-squared error RMSE and relative change in RMSE during the null-step trials for the early, mid, and late swing gait targets (ES, MS, and LS, respectively), applied during the Impaired Dynamics III condition. The visual dynamics cues (VCD) group (red) began practice seeing the virtual patient’s leg while the control group (blue) only saw the virtual patient’s ankle. *Significant group differences. Standard errors shown.

## DISCUSSION

In the context of a novel locomotor training task, i.e., training a virtual patient (VP) walking on a treadmill, this study tested the hypothesis that visual dynamics cues (VDCs) facilitate a human trainer’s ability to learn and manipulate locomotor dynamics. This was based on the rationale that seeing the geometry of the VP’s leg provides dynamics-relevant visual cues that may engender prior experiences with leg-like objects (i.e., linked segment systems influenced by gravitational, inertial, Coriolis, centrifugal, and in this case, muscular kinetics), which could help by informing and constraining the space of to-be-learned internal dynamics representations. The results suggest that VDCs did cause participants to invoke prior dynamics experiences; however, the hypothesis was not supported. VDCs failed to provide a learning benefit and in some cases (i.e., the late swing gait target), VDCs created a dependency that became apparent when the VP’s leg display was removed in a generalization task.

### Influence of Visual Dynamics Cues on Initial Dynamics Expectations

Showing the leg during the initial null-dynamics trials appeared to instantiate a specific set of expectations about what impedance forces would be felt by participants. Subjects who could see the VP’s leg (VDC group) lifted it too high during the swing phase, while those who only saw the ankle interaction point (control group) did not show this bias. It could be that upon seeing the leg, subjects expected to feel its weight and adjusted by lifting upwards to provide (unnecessary) support. This occurred even though subjects were explicitly told that, during the initial null-dynamics trials, they would not feel forces from the VP’s leg and they only had to trace through the air. This is consistent with evidence that explicit knowledge about an object’s properties (e.g., mass) supplied by non-proprioceptive modalities is not sufficient to override prior experiences, i.e., sensorimotor errors are needed to update internal models^19^.

An alternative explanation is that the large mid-swing target errors during the initial null-dynamics trials arose from the two-thirds power law, i.e., the inverse relation between movement curvature and speed^20^. After passing the early swing target, high curvature of the ankle trajectory is needed: the VP’s ankle must arc downward to hit the mid-swing target. Although the VP leg’s pendular dynamics would normally facilitate this motion, they were absent in the initial null-dynamics trials. Therefore, subjects may have rounded-off the VP’s ankle trajectory instead of slowing down. However, if this were true, it would be expected to affect both groups similarly, yet the control group was able to achieve the necessary curvature to reach the mid-swing target. Therefore, it is more likely that the high mid-swing error in subjects who initially saw the VP’s leg arose from their prior experiences, i.e., large body parts are usually heavy, brought about from the visual cues of the VP’s leg.

### Effects on Task Performance and Generalization

The results did not support the hypothesis that VDCs facilitate learning complex dynamics. Once the impaired locomotor dynamics were turned on (during Impaired Dynamics I), the subjects who could see the VP’s leg (VDC group) were able to achieve a level of manipulative accuracy close to that of participants who only saw the VP’s ankle (control group). Notably, group differences were observed for the generalization tests, which shifted the gait target positions. This required the manipulandum to be moved along a different path that changed the forces required for task success. To facilitate comparisons, both subject groups performed the generalization task under identical feedback conditions (end-point visual feedback only), meaning that the VDC group had to perform the task without seeing the VP’s leg. Although both groups performed similarly for the early and mid-swing targets, the VDC group had worse generalization for the late swing target, and only improved their accuracy in this condition after they were able to practice *without* seeing the VP’s leg for a period of time.

These generalization outcomes showing no VDC benefits contrast with those of a recent study by Sadeghi and colleagues^6^. In their study, subjects who practiced a task seeing the configuration of a virtual hammer (a circular handle with a mass on the end) had better generalization than those who used the same tool but saw an ambiguous display without the hammer head and handle. Using a computational model, the authors concluded that seeing VDCs improved subjects’ ability to transfer learning across different object actions (directions of hammer rotation) and that VDCs appeared to be a significant determinant of generalization relative to the information participants received from physical interaction (proprioception). These different study outcomes are likely because Sadeghi et al. tested participants under the same visual conditions, i.e., those who practiced while seeing the hammer geometry also saw it on generalization tests, and vice-versa. On the other hand, the present study used a mixed within/between subjects study design; the VDC group performed the generalization test *without* seeing the leg.

In the present study, the failure of VDCs to facilitate generalization can be most readily explained by the specificity of practice premise, i.e., retention and generalization are maximized in the same sensory context^21^. When the VP’s leg was visible during initial practice, participants may have used the geometry of the thigh and shank for error correction, leading to a dependency that became apparent when the subjects switched to practice a different generalization target arrangement without seeing the VP’s leg. This echoes the results of Proteau and colleagues^22^, who asked subjects to practice an aiming task and showed that when visual feedback of their own limbs was removed performance became worse in comparison to a control group who only saw end-point feedback. One could also speculate that the visual display of the leg influenced how participants internally represented the dynamic structure of the VP. If for example, the VDC group learned to represent the VP state in angular coordinates, an end-point transformation may have been needed to determine how much force to apply at the VP’s ankle (e.g., ***J***(***θ***)^−*T*^ [Eq. 3]; n.b., it is not implied that the nervous system uses this equation explicitly). However, when the VP leg segments were visually removed this learned transformation may no longer be helpful.

Finally, the observed group differences in the unexpected null step responses suggest that subjects in the VDC group may have incurred a “switching cost” in terms of adaptation. Specifically, the group who practiced seeing the VP’s leg had smaller null-step errors for the early swing target, which may reflect higher antagonistic muscular co-activation due to greater uncertainty in dynamics representations^23^. Note that in these types of catch trials, a larger error is thought to reflect stronger adaptation in a deterministic environment^24^. Thus, the null steps suggest less adaptation in the VDC group. Group differences in null-step responses were not observed for mid and late swing targets, which is likely because subjects may have had sufficient time to initiate corrective actions before reaching these targets. In this regard, force channels are a useful type of probe because they clamp kinematic errors and allow longer time scales to be investigated^25^. A previous study using the same VP gait training task implemented force channels by locking the VP’s ankle motion to a predefined target trajectory^11^. However, in the present study, force channels could not be used because there was no target trajectory, only discrete gait targets.

### Limitations

The results discussed above should be interpreted in the context of the study’s limitations: 1) Without further experiments, it is unclear whether the results would extend to different motor tasks, including the physical manipulation of a real person. 2) It is impossible to know what subjects were looking at without eye-tracking measurements, i.e., whether the VDC subjects foveated on the leg segments or something else. 3) Additional experiments and visual manipulations would be needed to make broader inferences about the effect of the visual display and other types of visual cues. 4) In this study the hypothesis did not test the specific mechanism(s) by which visual cues could act on internal representations (e.g., by restricting the choice of models, activating existing models, etc.); this would require a different study design and/or computational models of the human neural control system (e.g., see Sadeghi and colleagues^6^). 5) The mismatch in subject expectations during the initial null-dynamics trials for the VDC group could mask VDC benefits. Therefore, it is unknown whether VDC would facilitate learning if the VDC subjects started practice with impaired leg dynamics, which would be more congruent with subject expectations. Nevertheless, both subject groups had similar performance at the end of the initial null-dynamics practice block, which weighs against this possibility. 6) The extent to which the control group’s performance on the task depended on them knowing that they were interacting with a locomoting VP is unknown. The task instructions were the same for all subjects so that any observed differences would be due to the manipulation of the VDCs and not asymmetrical withholding of information.

### Implications

The present results add qualifiers to the conclusions of prior research showing that VDC enhances generalization in tool-use^6^. Generalization benefits may not extend to different sensory contexts and may erode as dynamics complexity increases (but note that further study is needed to firm these conclusions, e.g., by comparing sensory and dynamics manipulations within the same study population and task context). Ultimately, the findings convey an intuitive message in-line with the specificity-of-practice premise^26^: it is best to practice under the same visual conditions one would expect to encounter in real situations. If at some point VDCs may become unavailable, it may be better to practice without them, as there did not appear to be any learning deficit in the group that never saw the VP’s leg. This is well known by pilots, who often train for low- or null-visibility conditions (e.g., in clouds) using black-out hoods that obscure visual cues outside the cockpit. The results may also be of particular relevance in virtual and augmented reality, which can readily add or overlay visual information that is normally unavailable, for example through the use of interactive mirrors^27,28^.

## ACKNOWLEDGMENTS

Funded by Northeastern University and the National Center for Advancing Translational Sciences (NIH# UL1TR002544).

